# Neural tracking of stressed syllables in Dutch nursery rhymes relates to vocabulary outcomes in a large, longitudinal sample

**DOI:** 10.64898/2026.06.24.734253

**Authors:** Anika van der Klis, Katharina Menn, Melis Çetinçelik, Tineke M. Snijders, Caroline Junge

**Affiliations:** Experimental Psychology, Department of Psychology, Utrecht University, The Netherlands; Department of Cognitive Neuropsychology, Tilburg University, The Netherlands; Department of Cognitive Neuroscience, Maastricht University, The Netherlands; Max Planck Institute for Psycholinguistics, Nijmegen, the Netherlands

**Keywords:** neural tracking, speech-brain coherence, electroencephalography, brain development, vocabulary development, YOUth Cohort Study

## Abstract

Speech consists of regularities at different timescales. Already during infancy, neural electrophysiological activity aligns to these rhythms. The degree to which infants exhibit neural tracking of speech can be linked to their language development. In this study, we examined how the neural tracking of sung speech develops across age, from infancy to early childhood, and across different frequency bands (i.e., at the stress, syllabic, and phonemic rates), and whether neural tracking at each frequency and age predicts children’s language outcomes. We included 2565 children of the longitudinal YOUth cohort. Children listened to Dutch sung nursery rhymes while EEG was recorded at three measurement waves. After preprocessing the data, we included 955 children at 5 months, 1048 children at 10 months, and 795 children at 2-4 years. The final sample consisted of 750 children who also completed a receptive vocabulary test at 2-4 years. Children from 5 months onwards showed significant neural tracking of stressed syllables, syllables, and phonemes, measured with speech-brain coherence (SBC). Unexpectedly, there were no developmental changes in SBC across different frequency bands from infancy to early childhood. As expected, children with larger receptive vocabularies showed increased SBC in the stressed syllable rate. These findings suggest that stronger tracking of stressed syllables is related to individual differences in language ability.

## 1. Introduction

Infants, even in the first months of life, can make sense of the complex and dynamic world around them. From a constant stream of incoming information – like speech – they can extract structure, detect patterns, and start to understand meaning (Gervain & Mehler, 2010; Romberg & Saffran, 2010; Swingley, 2009). The rhythmic properties of infant-directed speech are predictable and help the listener to segment the continuous speech stream into smaller meaningful units (Johnson & Jusczyk, 2001; Jusczyk et al., 1999; Snijders & Menn, 2026). In a classic experiment, Jusczyk et al. (1999) showed that 7.5-month-olds can use syllable stress to segment bisyllabic words from continuous speech. There is also neural evidence for infants’ sensitivity to rhythmic properties of speech. Studies using electroencephalography (EEG) show that when listening to continuous speech, the infant EEG signal aligns with prosodic stress in both spoken and sung speech (Attaheri, Choisdealbha, et al., 2022; Kalashnikova et al., 2018; Menn, Michel, et al., 2022) which has been related to behavioural outcomes of word learning (Attaheri et al., 2024; Menn, Ward, et al., 2022; Snijders, 2020). This alignment between rhythmic properties of speech and electrophysiological activity is referred to as neural tracking. It has been hypothesised that it may be based on the synchronisation of oscillatory brain rhythms with rhythmic units in speech (Giraud & Poeppel, 2012; Goswami, 2018).

Neural oscillations provide temporal windows of enhanced excitability in neuronal networks (Buzsáki & Watson, 2012), making neurons more sensitive to input during moments of higher excitability. Therefore, the temporal alignment of neural oscillations to linguistic elements may help the brain to process relevant information while suppressing any distracting input. The brain generates rhythmic oscillations at varying frequencies. At rest, neural oscillations in the auditory cortex are hierarchically organised, including the delta (1-3 Hz), theta (4-8 Hz), and low gamma (25-35 Hz) frequency ranges which approximately correspond to phrasal or word stress, syllabic, and phonemic time scales in speech, respectively (Giraud & Poeppel, 2012). Currently, it has been shown that electrophysiological activity synchronises with the rate of phonemes and phonetic features (Di Liberto et al., 2015, 2023; Menn et al., 2023b), syllables (Ding et al., 2016; Peelle et al., 2013; Pefkou et al., 2017), and intonation phrases (Ding et al., 2016; ten Oever et al., 2022). It has been suggested that this alignment between electrophysiological brain activity and speech information is an important mechanism underlying speech processing, as it may help the listener to segment continuous speech into linguistically meaningful units, e.g., phonemes, words, and phrases (Meyer, 2018).

In adults, neural tracking of speech is shaped by the acoustic properties of the speech signal, linguistic knowledge, and attention. Manipulations of the speech signal, such as increasing the speech rate or introducing noise, impacts the degree of neural tracking (Pefkou et al., 2017; Verschueren et al., 2022). In addition to bottom-up processing of the acoustic signal, linguistic knowledge and attention modulate the neural tracking of speech. For example, Ding et al. (2016) presented Chinese stimuli that did not contain acoustic cues marking sentence boundaries to Chinese- and English-speaking participants. Only Chinese-speaking participants showed responses to the phrasal rhythm. In multispeaker “cocktail party” environments, where listeners have to filter out competing conversations, listeners show neural tracking of both attended and unattended speech, but tracking of attended speech is stronger and associated with improved comprehension (Barchet et al., 2025; Zion Golumbic et al., 2013). These findings indicate that adult neural tracking reflects an interaction between acoustic features of the speech signal, linguistic processing, and selective attention.

In infants and children, neural tracking is similarly influenced by acoustic features, linguistic knowledge, and emerging attentional processes. For example, infants show enhanced neural tracking of infant-directed speech compared to adult-directed speech, likely due to its exaggerated amplitude modulations and prosodic structure or increased attention to this speech register (Kalashnikova et al., 2018; Leong et al., 2017; Menn, Michel, et al., 2022). Familiarity or top-down knowledge also affects neural tracking already early in life: Newborns exhibit increased neural tracking when listening to their native compared to a foreign language, but only when the foreign language differs rhythmically, suggesting an early sensitivity to familiar rhythmic patterns (Ortiz-Barajas et al., 2023). In childhood, background noise can disrupt neural tracking. For instance, van Hirtum et al. (2023) showed that in 5-year-olds, theta-band tracking was drastically reduced for speech in noise, while delta-band tracking was related to behavioural measures of speech intelligibility. These developmental findings suggest that while infants and children are sensitive to bottom-up acoustic cues, the role of top-down factors such as attention continues to mature.

Research on neural tracking across development reveals a complex and sometimes inconsistent pattern of age-related changes from infancy through adulthood. Regarding infants, Ortiz Barajas et al. (2021) compared the phase and envelope tracking in newborns to 6-month-olds to examine whether language familiarity influences the degree of neural tracking. They found that infants track the phase of the familiar and unfamiliar language regardless of age, while the amplitude was only tracked by newborns and by 6-month-olds in the unfamiliar language. In a longitudinal study, Attaheri, Choisdealbha, et al. (2022) examined neural tracking in 60 infants at 4, 7, and 11 months of age while they listened to nursery rhymes. The presence and strength of neural tracking was assessed in three frequency bands: delta (0.5-4 Hz), theta (4-8 Hz), and alpha (8-12 Hz) for control. The authors found a significant effect of age on neural tracking in the delta frequency, where 4-month-olds show increased neural tracking compared to 11-month-olds. For young children, Rogachev and Sysoeva (2024) cross-sectionally found an increase in neural tracking between 3 and 8 years of age. In contrast, Bertels et al. (2023) examined neural tracking between 5 and 27 years of age using magnetoencephalography (MEG). For speech without noise, they found no effect of age on phrasal tracking. For syllabic tracking, they only found an increase in the right hemisphere between 7.5 and 10.5 years of age. Lastly, Attaheri, Panayiotou, et al. (2022) compared their infant sample to adults. Using the same paradigm, the authors did not find many differences in neural tracking of sung speech between infants and adults, although the data show a trend of age-related increases in the faster alpha and theta bands. This pattern can be predicted by maturation of the brain, as there is a general speed-up and increase in activity in faster frequencies across early childhood (Anderson & Perone, 2018; Menn et al., 2023a; Snijders & Menn, 2026). Taken together, these findings indicate that different aspects of neural tracking are differentially modulated by age. Although the findings remain inconclusive, most studies suggest that neural tracking reflects subtle frequency-specific and context-dependent changes that possibly align with broader patterns of brain maturation.

The degree to which infants and children exhibit neural tracking of speech can be linked to both their concurrent and later language abilities (Attaheri et al., 2024; Çetinçelik et al., 2023, 2024; Menn, Ward, et al., 2022; Nguyen et al., 2023; Rogachev & Sysoeva, 2024). For example, cross-sectional research demonstrates that the neural tracking of stressed syllables in nursery rhymes at 10 months, but not at 14 months, predicts vocabulary outcomes at 24 months (Menn, Ward, et al., 2022). In line with this, Attaheri et al. (2024) found that 11-month-olds’ delta band tracking predicts vocabulary at 24 months. In addition, it was found that the neural tracking of stress and syllables at 10 months significantly predicted vocabulary at 18 months (Çetinçelik et al., 2023, 2024). Thus, ample evidence shows that the neural tracking of low-frequency information in late infancy can predict future language skills. It should be mentioned that these studies used different stimuli (sung versus spoken speech), and the syllable rate in the infant-directed speech used in Çetinçelik et al. (2023) fell under the delta frequency rate, which was the significant predictor of vocabulary in Attaheri et al. (2024). It remains unclear whether the relation with language is specific for neural tracking in the delta frequency band or whether it is specifically related to the neural reaction to specific stimulus characteristics (Snijders & Menn, 2026). Although most evidence is based on infants, Rogachev and Sysoeva (2024) examined children aged 3-8 years and found a positive association between processing of both acoustic and semantic information with children’s concurrent receptive language skills.

Together, these studies provide evidence that children’s neural tracking of speech is related to their language development (Cantiani et al., 2022). For vocabulary development, the tracking of low-frequency information could be particularly relevant (Attaheri et al., 2024; Çetinçelik et al., 2024; Menn, Ward, et al., 2022). For stress-timed languages, such as Dutch, initial word stress is an important cue for continuous speech segmentation (Cutler, 1994). Indeed, experimental studies have shown that infants use such rhythmic properties of speech to segment words from the speech stream (Cutler & Butterfield, 1992; Jusczyk et al., 1992; Kooijman et al., 2009). Therefore, children attending to word stress in the speech stream might benefit in the word segmentation task, subsequently facilitating language development (Junge et al., 2012; Snijders, 2020).

There is some evidence suggesting that infants’ ability to neurally track speech decreases across the first year of life, at least in the word stress rate and when examining amplitude tracking (Attaheri, Choisdealbha, et al., 2022; Ortiz Barajas et al., 2021). Comparing their infant to adult sample, Attaheri, Panayiotou, et al. (2022) showed that there were only minimal changes in neural tracking between infants and adults in each frequency rate. However, the data pointed towards a trend of stronger neural tracking in higher frequencies, including theta and alpha, for adults. It is still unknown how the neural tracking of different frequencies within speech develops from infancy until early childhood. Therefore, the first research question of the current study is: “How does neural tracking develop across different frequency rates from infancy to early childhood?”. To examine this, we collected continuous EEG while children listened to Dutch nursery rhymes at three different ages: 5 months; 10 months; and 2-4 years. Our hypothesis is that, due to brain maturation, neural tracking gets stronger for faster relative to slower frequencies across early development (Menn et al., 2023a; Snijders & Menn, 2026). This is the first study to examine neural tracking in such a large sample of neurotypically developing children. Across development, we expect a decrease in neural tracking of the stressed syllable rate (Attaheri, Panayiotou, et al., 2022), and possibly an increase in neural tracking of the faster frequency rates (Menn et al., 2023a; Snijders & Menn, 2026).

Previous studies with infants found that increased neural tracking in the delta band (i.e., overlapping with stress rate and slow syllable rate), but not the theta band (i.e., overlapping with syllabic rate), predicted later vocabulary (Attaheri et al., 2024; Çetinçelik et al., 2023; Menn, Ward, et al., 2022). Menn, Ward, et al. (2022) conducted a cross-sectional study in which they found that neural tracking of the stress rate of sung nursery rhymes at 10 months (but not at 14 months) can predict children’s vocabulary outcomes at 24 months. The second research question of the current study is: “Do individual differences in neural tracking at specific rates and ages predict children’s vocabulary outcomes?”. Based on previous research using the same sung nursery rhymes (Menn, Ward, et al., 2022), we expect that infants’ neural tracking in the stress rate at 10 months is especially relevant for children’s vocabulary outcomes, because around 10 months Dutch infants are using stress patterns to learn to segment words from continuous speech (Cutler & Butterfield, 1992; Kooijman et al., 2009; Vroomen et al., 1996). To examine this, we collected children’s receptive vocabulary scores using the Peabody Picture Vocabulary Test (PPVT-III-NL) at 2-4 years. Although we did not preregister specific hypotheses for tracking at 5 months and 2-4 years, we envisage several possible outcomes. At 5 months, neural tracking of the stress or syllabic rate can relate to later language skills, as previous research suggests even young children may already be able to segment words based on word initial stress (Höhle et al., 2009; Thiessen & Erickson, 2013) or transitional probabilities between syllables (Johnson & Jusczyk, 2001; Thiessen & Erickson, 2013; Thiessen & Saffran, 2003). At 2-4 years, we hypothesise that neural tracking of the acoustic rhythms might not relate to children’s vocabulary skills. At this age, top-down linguistic knowledge is more important for speech processing, making low-level acoustic cues potentially less important. The results of the current study will shed more light on the role of neural tracking of (sung) speech in children’s language development.

## 2. Materials and methods

### 2.1. Participants

The data for this study are collected as part of YOUth, a longitudinal cohort study part of Utrecht University and University Medical Center Utrecht (Onland-Moret et al., 2020). For this study, we started with the total sample of 2565 participants. Children participated in various tasks at regular intervals (“waves”), including the EEG task analysed for this study. At 5 months, EEG data were collected for 1851 participants. At 10 months, EEG data were collected for 1650 participants. At 2-4 years, EEG data were collected for 979 participants. After preprocessing the data (see section 2.3), we could include 1757 participants with at least 70 clean epochs in at least one session. At 5 months, there are data of 955 children (*M* = 5.5 months; *SD* = 0.9 months; 50% female), 1048 children at 10 months (*M* = 10.4 months; *SD* = 1.1 months; 48% female), and 795 children at 2-4 years (*M* = 3.7 years; *SD* = 0.8; 49.4% female). There are 906 children that have EEG data left in only one of the waves, 661 children who have EEG data left in two waves, and 190 children who have EEG data left in all three waves.

Of this preprocessed sample, 933 children aged ≥2 years and 3 months (i.e., within the normative range for scoring) participated in the receptive vocabulary test in the lab at Wave 3. From the analyses on language outcomes, we excluded children as preregistered on OSF: [link removed for review]. First, we excluded all children who were multilingual defined as hearing less than 80% Dutch at home (*n* = 145). We collected a questionnaire around birth, based on which we excluded all children who were born preterm (<= 36 weeks) and have a low birthweight (<= 2500 gram) (*n* = 40). We also excluded all children with hearing problems at birth (*n* = 51) or a developmental disorder reported by caregivers at 2-4 years (*n* = 39). Lastly, we excluded all children who did not fully finish the receptive vocabulary task in the lab (*n* = 76). The final sample consisted of 750 infants included in the reported analyses concerning the relationship with language development, fully described across the three waves in Table 1.

**Table 1:** Descriptive characteristics of the final analytic sample (preprocessed EEG inclusion with valid vocabulary score) across three measurement waves.

| Variable | Wave 1 (5 months)<br>(*n* = 357) | Wave 2 (10 months)<br>(*n* = 402) | Wave 3 (2-4 years)<br>(*n* = 592) |
| --- | --- | --- | --- |
| Age in weeks, *M* (*SD*) | 23.6 (3.6) | 45.4 (4.6) | 194.3 (39.7) |
| Gender (%) |  |  |  |
| • Female | 174 (48.7%) | 204 (50.8%) | 292 (49.3%) |
| • Male | 183 (51.3%) | 198 (49.2%) | 300 (50.7%) |
| *M* (*SD*) reported Dutch exposure (%) | 98.5 (3.0) | 98.6 (2.7) | 98.5 (3.2) |

*Note.* *M* = Mean. *SD* = Standard Deviation.

The YOUth cohort study is carried out in accordance with The Code of Ethics of the World Medical Association (Declaration of Helsinki). All caregivers have signed informed consent before data collection. The YOUth baby cohort framework protocol was approved by the medical ethical committee of the University Medical Center Utrecht (application number 14-616). Infants received a romper with the YOUth cohort logo at Wave 1, a Miffy picture book after participating in Wave 2, and a frog umbrella after participating in Wave 3. Caregivers received a small monetary compensation after each session (€30).

### 2.2. Materials and procedure

#### 2.2.1. EEG task

Children sat on their caregivers’ lap or in a highchair at approximately 65 cm distance from the screen in a dimly lit and quiet room. The labs were semi-dark and controlled for luminance. Continuous EEG was recorded using a 32-channel ActiveTwo BioSemi system collecting data at a 2048Hz sampling rate. Recordings were made from 16 left and 16 right hemisphere electrodes positioned according to the international 10-20 system. The Common Mode Sense (CMS) and Driven Right Leg (DRL) circuits were used to reduce common-mode interference. During the EEG study, children were presented with different stimuli, of which only one video of sung nursery rhymes is relevant for the current analysis. The sung nursery rhyme video consisted of two women singing five Dutch nursery rhymes that are familiar to Dutch infants (translated from Jones et al., 2015; used in Menn, Ward, et al., 2022) (“Dit zijn mijn wangetjes” (These are my cheeks); “De wielen van de bus” (Wheels on the bus); “Hansje pansje kevertje” (Hansje pansje beetle); “Twinkel twinkel kleine ster” (Twinkle twinkle little star); and “Papegaaitje leef je nog?” (Parrot are you still alive?)). The women sung in an infant-directed way (such as high intonation, wide intonation range) accompanied by typical infant-directed behaviours (such as positive affect, idiosyncratic gestures). The two women alternately sung one of the five nursery rhymes, with the total duration for the five songs being 69 seconds. The nursery rhyme video was alternated with a video of colourful, moving toys (not analysed for the current report), with both videos repeating three times. Therefore, the total duration of analysed data in one session was 3 * 69 = 207 seconds. For a previous study (Menn, Ward, et al., 2022), the duration of all stressed syllables, syllables, and phonemes were transcribed using Praat (Boersma & Weenink, 2020). In the videos, 85% of stressed syllables occurred at a rate of 1-3 Hz and 85% of all phonemes occurred at 5-15 Hz. The non-overlapping rate of 3-5 Hz mostly captured syllables.

#### 2.2.2. Vocabulary task

During Wave 3 at 2-4 years, we administered the third version of the Dutch Peabody Picture Vocabulary Test (PPVT-III-NL; Schlichting, 2005). It is a multiple-choice test that measures whether children can match one of four pictures to the correct audio stimuli, therefore measuring children’s receptive vocabulary (i.e., language comprehension). We used a computerised version where pre-recorded audio stimuli were played on a computer, and children pressed on one of the four pictures presented on the touchscreen. Items become increasingly more complex when the child progresses further in the task. The task terminates when participants make nine or more errors in one set of words consisting of 12 items. The program automatically calculated children’s raw scores by subtracting the number of errors from the total number of presented items (final set * 12 items). Normed scores (*M* = 100; *SD* = 15) were determined from the raw scores using the child’s age during testing and the tables provided in the official handbook (Schlichting, 2005). Caregivers could be present in the room, but they were seated in the back out of the child’s view and explicitly instructed not to help or communicate with the child about the task.

### 2.3. EEG preprocessing

EEG data of all 32 electrodes were analysed in Matlab using a fully automated custom pipeline that used the FieldTrip toolbox (Oostenveld et al., 2024), EEGLab (Delorme & Makeig, 2004), and iMARA (Marriott Haresign et al., 2021). This preprocessing pipeline was selected as it gave the best test-retest reliability in a separate sample of 10-month-olds (Chaudhry et al., 2025). Recordings were first down sampled to 512 Hz, re-referenced to Cz, and filtered using the Butterworth filter (high-pass filter = 0.1 Hz, low-pass filter = 40 Hz, notch filter = 50 Hz). For preprocessing only, the data were segmented into 1s epochs and checked for the following criteria to detect noisy channels: absolute amplitude > 250 γV, flatlining 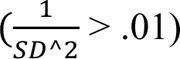, kurtosis > 10, and variance > 2000 μ V^2^ (*SD*^2^). If more than 40% of the segments in a channel exceeded any of these limits, the channel was flagged as noisy. If the session had more than 3 noisy channels, the session was removed from subsequent preprocessing steps. After these initial preprocessing steps, continuous data of the remaining channels were cleaned using Independent Component Analysis (ICA) using *runica* of EEGLab and iMARA for selecting artefactual components. The noisy channels were interpolated using spherical splines. Lastly, the remaining data were re-referenced to the common average of all electrodes. After that, the data were segmented into 3s epochs using a 1s sliding window. To detect artefactual epochs, the same criteria as for noisy channel detection were used. Only children with at least 70 clean epochs were included in the speech-brain coherence (SBC) analyses (*n* = 1757).

### 2.4. Analyses

#### 2.4.1. Speech-brain coherence

Following Menn et al. (2022), we first computed the speech envelope of the stimuli using a Hilbert transform with a 4^th^-order Butterworth filter. We then used a Fourier transform of both the speech envelope and the EEG signal from 1-15 Hz. Subsequently, SBC was calculated as the cross-spectrum between the EEG signal and the speech signal normalised by the power spectra of these signals. Coherence reflects the consistency of the phase difference between the two signals at a given frequency:

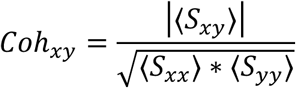

We also created surrogate data by randomly shuffling the speech envelope across epochs and then computing SBC.

SBC values were normalised by the surrogate data based on the same number of trials to ensure the numbers of trials did not affect the findings (see Bastos & Schoffelen, 2015). We used the following formula following Menn et al. (2022):

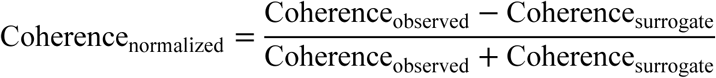

Noteworthy, these calculations are an exact replication of the approach in Menn, Ward, et al. (2022), using the exact same stimuli on a different longitudinal sample.

SBC for observed and surrogate data was averaged across channels and across three different frequency bands: stress range (1-3 Hz), syllabic range (3-5 Hz), and phonemic range (5-15 Hz). We used *t*-tests for each frequency band (stress; syllabic; phonemic) to examine whether the mean of observed data was significantly higher than the mean of surrogate data, indicating that children tracked the speech signal above chance-level at that given frequency.

#### 2.4.2. Statistical analyses

We fitted linear mixed-effects regression models using the *lme4* package (version 1.1-35.2; Bates et al., 2015) and the *lmerTest* package to obtain *p*-values (version 3.1-3; Kuznetsova et al., 2017) in *R* 4.3.3 (R Core Team, 2025). The continuous predictors (age and vocabulary) were centred and scaled in all models. For the Frequency variable, we used treatment (dummy) coding with Phonemic as the reference category.

The wide age range within waves can make it more difficult to detect any non-linear age effects that may occur within measurement waves, particularly at Wave 3 (i.e., 2-4 years). Therefore, we first fitted multiple linear mixed-effects regression models separately for each measurement wave to examine the effects of age (linear) and age^2^ (quadratic) separately within these age groups, using the following formula: SBC ∼ age + age^2^ + (1|participant). These models show whether there are any significant effects of linear age or quadratic age on SBC within measurement waves.

Then, on the pooled dataset, we fitted the following model: SBC ∼ age * frequency * vocabulary + (1|participant). We first assessed whether adding age^2^ to this model significantly improved model fit using the anova() function. The results first reveal whether there are any significant effects of linear age or quadratic age on SBC which answers the first research question. If age^2^ does not significantly improve the fit of the full model, we report the simpler model without age^2^.

To answer the second research question, we examined whether vocabulary or any of the interactions with vocabulary has a significant effect on SBC in the full model. Any significant interactions can reveal whether the relation between vocabulary size and SBC is dependent on children’s age or the frequency rate.

## 3. Results

### 3.1. Speech-brain coherence across early childhood

The descriptives for the standardised SBC values per wave and frequency band are depicted in Figure 1. At each frequency rate, SBC was significantly higher for the observed data than for the surrogate data. Welch’s t-tests showed significant differences between observed data and surrogate data in the phonemic range, *t*(5533.4) = 51.13, *p* < .001, syllabic range, *t*(5405.7) = 52.45, *p* < .001, and stress range, *t*(5265.2) = 74.02, *p* < .001.

**Figure 1:**
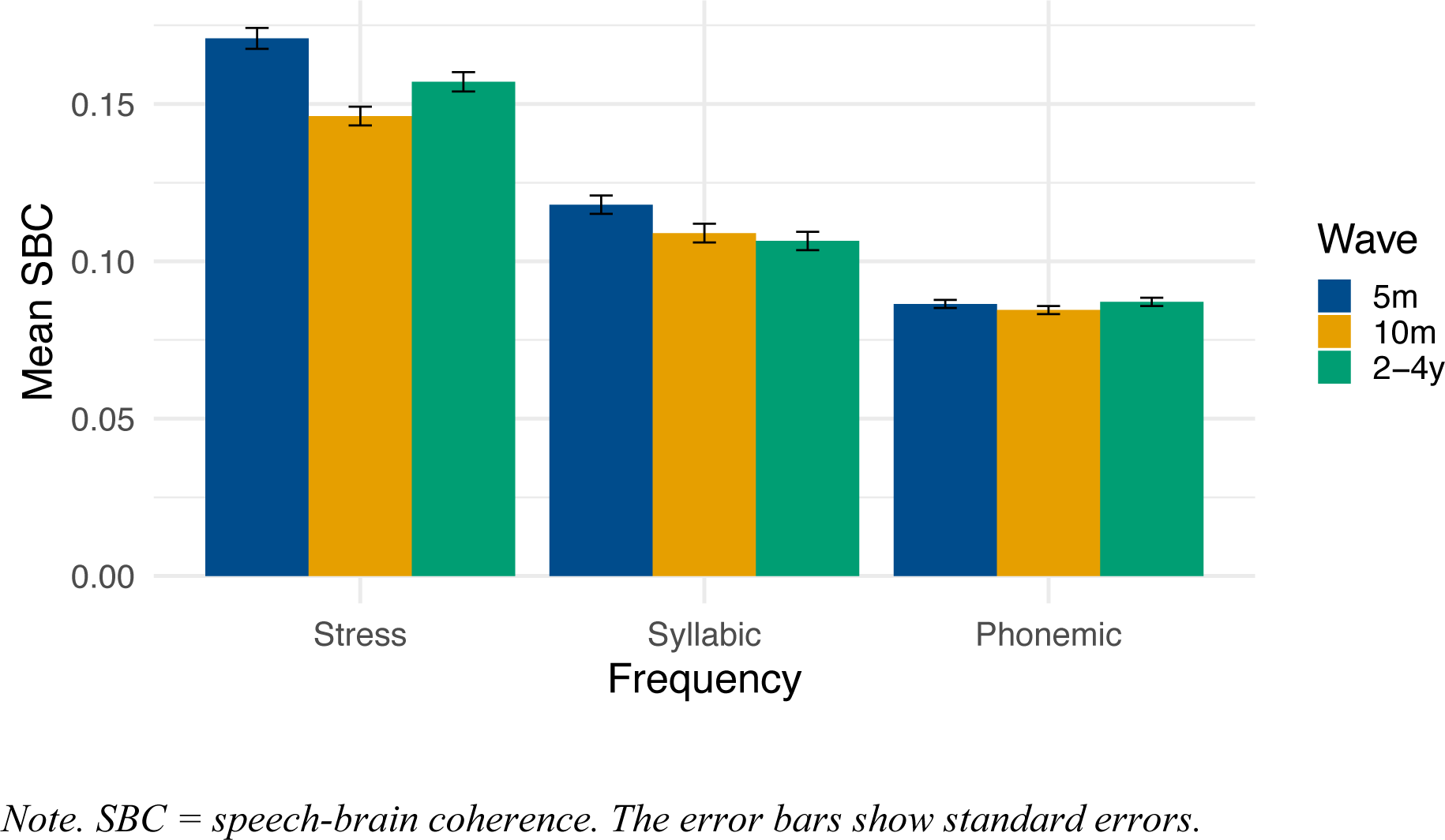
Average standardised speech-brain coherence in each frequency range per measurement wave

### 3.2. Development of speech-brain coherence

Given the wide age range within sessions, we first fitted three models assessing whether adding linear age and a quadratic age term (age^2^) on SBC significantly improved model fit for each wave separately. Figure 2 shows the developmental trajectories of SBC in each frequency rate within each measurement wave. Mixed-effects models with random intercepts for participant were initially fitted for all analyses. The models for Wave 1 (5 months) and Wave 3 (2-4 years) showed singular fits indicating negligible random-effect variance. Instead, equivalent ordinary least squares models were used for these waves.

**Figure 2:**
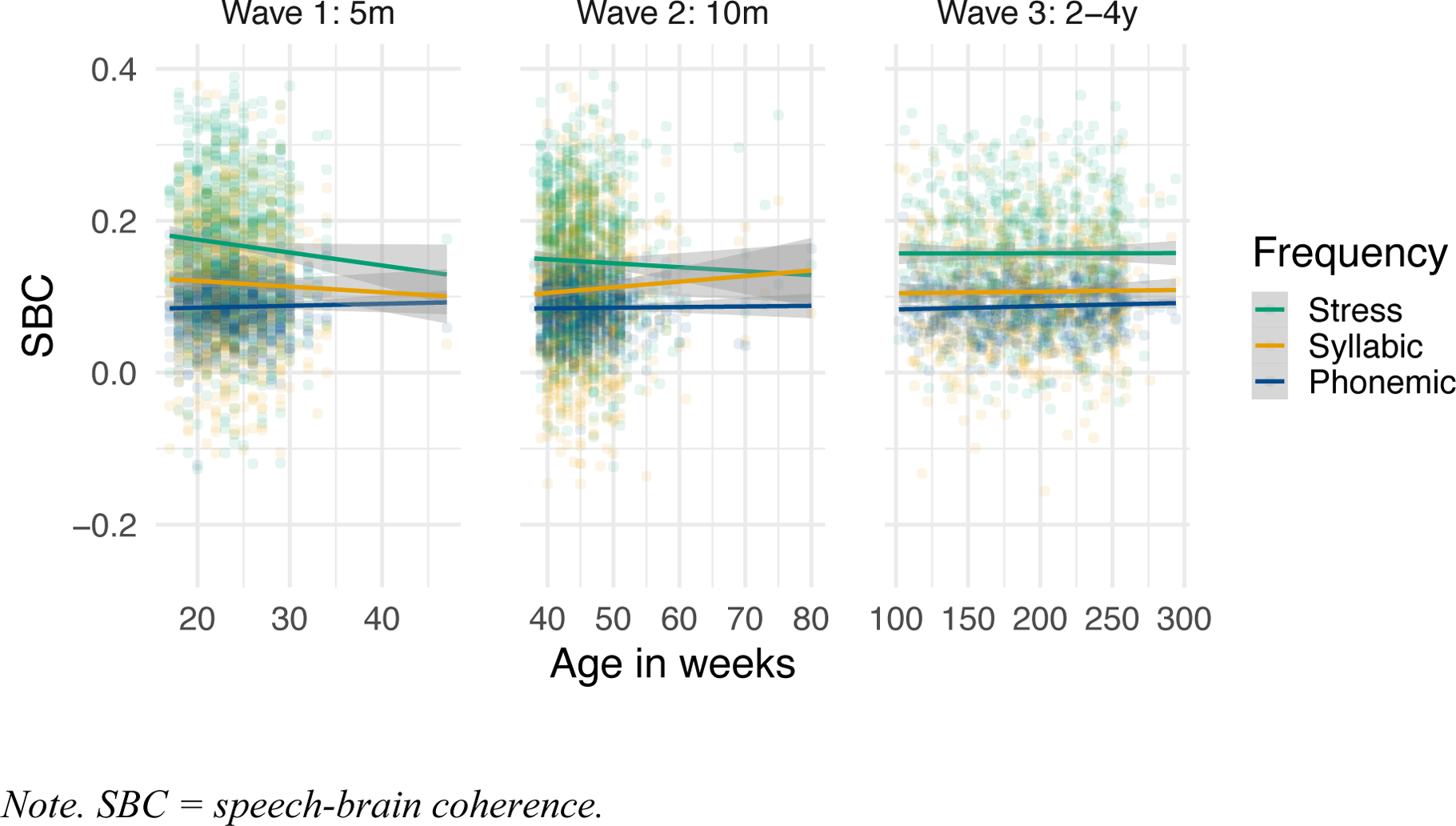
A scatterplot showing the distribution of normalized speech-brain coherence across the three frequency rates and measurement waves

At 5 months, a sequential ANOVA showed that adding linear age to the null model did not significantly improve model fit, *F*(1, 2461) = 3.38, *p* = .066. Adding age^2^ (squared) also did not significantly improve model fit compared to the model including linear age, *F*(1, 2460) = 0.55, *p* = .460) and compared to the null model, *F*(2, 2460) = 1.96, *p* = .141. At 10 months, a likelihood-ratio test comparing linear mixed-effects models with random intercepts for participant indicated that adding linear age did not significantly improve model fit relative to the null model, χ²(1) = 0.001, *p* = .973. Adding a quadratic age term also did not significantly improve model fit over the model including linear age, χ²(1) = 1.81, *p* = .179, and compared to the null model, χ²(2) = 1.81, *p* = .405. At 2-4 years, a sequential ANOVA showed that adding linear age, *F*(1, 1912) = 0.36, *p* = .547, or the quadratic age term, *F*(2, 1911) = 1.07, *p* = .342, as predictors did not significantly improve model fit compared to the null model. Adding the quadratic age term to the linear age model also did not significantly improve model fit, *F*(1, 1911) = 1.79, *p* = .182. Thus, across the three waves, the null model is favoured in all cases, indicating that there are no linear or quadratic effects of age on SBC within any of the measurement waves.

### 3.3. Relations to vocabulary outcomes

Next, we examined whether there were any significant effects of age, age squared, frequency (stress, syllabic, phonemic) and children’s receptive vocabulary skills (measured using the PPVT-III-NL) on SBC across all age groups. We fitted this model to all data pooled together, while adding a random intercept for participant. Adding age squared to the full model did not significantly improve model fit, χ²(1) = 3.50, *p* = .061, so we report the simpler model without age^2^.

The results in Table 2 showed a significant effect of Frequency, where SBC is significantly higher in both the stress (*b* = 0.069, 95% CI [0.064, 0.075], *p* < .001) and syllabic (*b* = 0.023, 95% CI [0.018, 0.028], *p* < .001) rates compared to the phonemic rate. Indeed, the highest mean was observed in stressed syllable tracking (*M* = 0.156, 95% CI [0.152, 0.159]), followed by syllabic tracking (*M* = 0.108, 95% CI [0.105, 0.112]), and phonemic tracking (*M* = 0.086, 95% CI [0.083, 0.090]) (see Figure 1). Tukey-adjusted pairwise comparisons indicated that all pairwise differences were statistically different (all *p*s < .001). These differences were not influenced by the age of children.

**Table 2:** Results of the linear mixed-effects regression model predicting speech-brain coherence using Age, Frequency, Vocabulary, and possible two-way and three-way interactions.

|  | *B [95% CI]* | Std.<br>Error | *t*-value | *p*-value |
| --- | --- | --- | --- | --- |
| (Intercept) | 0.086<br>[0.082, 0.090] | 0.002 | 42.105 | <.001*** |
| Age | 0.001<br>[-0.003, 0.004] | 0.002 | 0.421 | 0.674 |
| **Frequency:Stress** | **0.069**<br>**[0.064, 0.075]** | **0.003** | **24.778** | **<.001\*\*\*** |
| **Frequency:Syllabic** | **0.023**<br>**[0.018, 0.028]** | **0.003** | **8.205** | **<.001\*\*\*** |
| Vocabulary | -0.001<br>[-0.005, 0.003] | 0.002 | -0.677 | 0.499 |
| Age * Frequency:Stress | -0.001<br>[-0.006, 0.004] | 0.002 | -0.403 | 0.687 |
| Age * Frequency:Syllabic | -0.003<br>[-0.007, 0.002] | 0.002 | -1.176 | 0.240 |
| Age * Vocabulary | 0.001<br>[-0.002, 0.005] | 0.002 | 0.730 | 0.465 |
| **Frequency:Stress \***<br>**Vocabulary** | **0.005**<br>**[0.000, 0.011]** | **0.003** | **1.966** | **0.049\*** |
| Frequency:Syllabic *<br>Vocabulary | 0.003<br>[-0.002, 0.009] | 0.003 | 1.122 | 0.262 |
| Age * Frequency:Stress *<br>Vocabulary | 0.001<br>[-0.004, 0.005] | 0.002 | 0.303 | 0.762 |
| Age * Frequency:Syllabic *<br>Vocabulary | 0.000<br>[-0.005, 0.004] | 0.002 | -0.082 | 0.934 |
| Number of observations | 4053 |  |  |  |
| Conditional $R^2$ | 0.17 | | | |
| Marginal $R^2$ | 0.15 | | | |

*Note.* \* $p < .05$ ; \*\* $p < .01$ ; \*\*\* $p < .001$

We did not find a main effect of Vocabulary on SBC while the reference level was on the Phonemic rate (*b* = −0.001, 95% CI [−0.005, 0.003], *p* = .499). There was a small but significant interaction between Frequency (Stress) and Vocabulary, with children’s receptive vocabulary being stronger related to SBC in the stress rate compared to the phonemic rate (*b* = 0.005, 95% CI [0.000, 0.011], *p* = .049; see Table 2). The interaction is depicted in Figure 3 using the CRAN package *interactions* (Long, 2019). As shown in Figure 3, although there were no relationships between SBC and children’s receptive vocabulary size in the phonemic or syllabic range, we did find a significant positive increase in the stress range related to increases in receptive vocabulary. This interaction was not modulated by Age in a three-way interaction (*b* = 0. 002, 95% CI [−0.004, 0.005], *p* = .762; see Figure 4).

**Figure 3:**
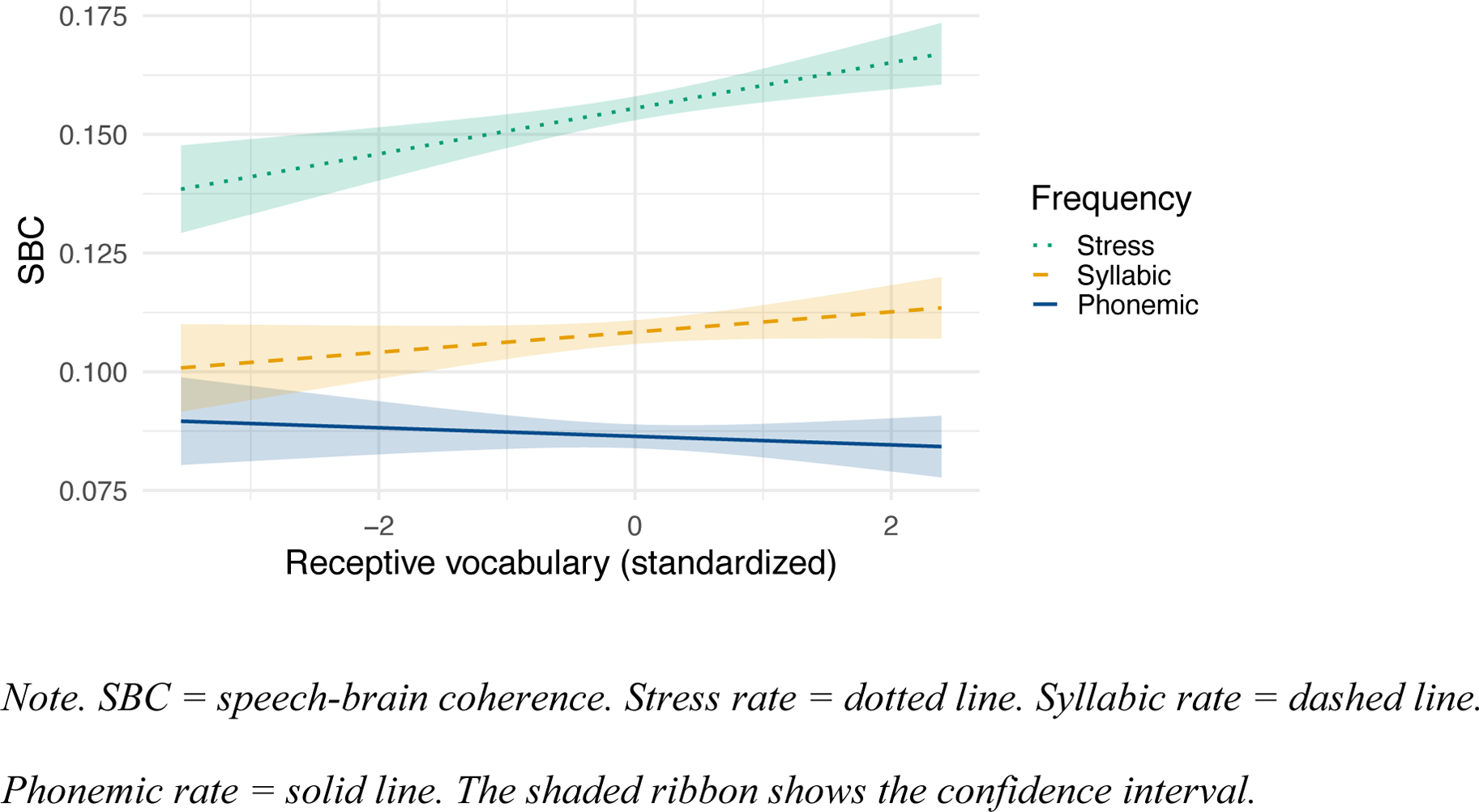
The interaction effect between Frequency and Vocabulary on speech-brain coherence

**Figure 4:**
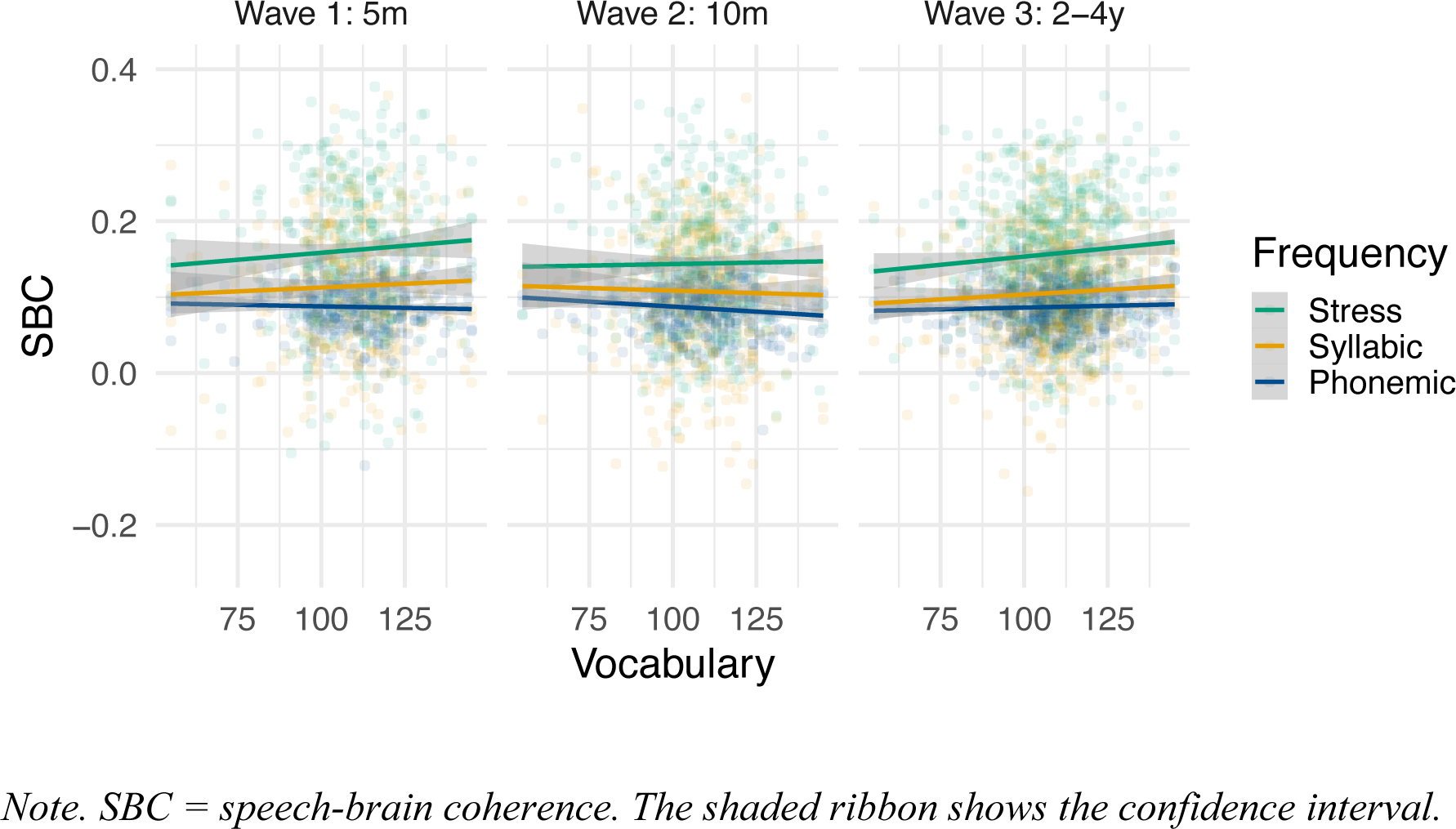
The association between children’s normed receptive vocabulary scores and speech-brain coherence across the three frequency rates and measurement wave

To probe the significant interaction between Frequency (Stress) and Vocabulary, we reparameterized the model with Frequency (Stress) as the reference level, allowing us to estimate the simple slope of Vocabulary on SBC in the Stress rate. In this parameterization, Vocabulary significantly predicted SBC (*b* = 0.004, 95% CI [0.000, 0.008], *p* = .044). The full mixed-effects model is reported in the *R* markdown script on OSF.

## 4. Discussion

In this study, we examined the longitudinal development of the neural tracking of Dutch sung nursery rhymes, measured with SBC, from infancy to early childhood. Although we did not expect any large developmental changes in neural tracking, we did expect that the relative tracking of faster frequencies (i.e., in the syllabic and phonemic rate) would increase across development due to brain maturation. In addition, we expected that children’s neural tracking of the stress rate would be positively associated with children’s language skills. This is the first study to address these questions in such a large, longitudinal cohort study, including data of 2565 children, contributing a total of 4480 longitudinal EEG-recordings of typically developing children measured during infancy and early childhood.

First, we found that 5-month-old infants, 10-month-old infants, and 2–4-year-old preschoolers neurally track Dutch nursery rhymes in the stressed syllable, syllabic, and phonemic rates significantly above chance-level. This matches the results of earlier studies with newborns and infants (e.g., Attaheri, Choisdealbha, et al., 2022; Ortiz Barajas et al., 2021). Observed SBC was significantly higher than surrogate SBC based on the same number of included epochs. Although we used a different method to measure neural tracking (SBC instead of temporal response function (TRF)), we found the same relative strengths of coherence across three frequency bands as Attaheri et al. (2022), with strongest coherence in the stress rate, weaker coherence in the syllabic rate, and weakest coherence in the phonemic rate. All these frequency differences were statistically significant. Therefore, the results of our study show that infants and preschoolers show strongest neural tracking in the slowest stress rate and weaker tracking in the faster frequencies, including syllabic and phonemic rates. The increase in SBC in the stress rate compared to the other rates is possibly amplified by the speech register used when recording the nursery rhymes used in this study. Although infants also show stronger neural tracking of stressed syllables compared to syllables in adult-directed speech, Menn, Michel, et al. (2022) showed that infant-directed speech was associated with an increase in infants’ SBC in the stress rate particularly. In line with this, our data show that, in nursery rhymes sung in an infant-directed manner, infants and preschoolers show the strongest neural tracking of stressed syllables.

Contrary to our predictions, we did not find developmental changes in children’s neural tracking of the stressed syllable, syllabic, and phonemic rates between 5 months and 2-4 years. We expected that there would be a decrease in stressed syllable tracking during infancy, as was previously found by Attaheri, Choisdealbha, et al. (2022) between 4 and 11 months, and a relative increase in the neural tracking of faster frequencies, including syllables and phonemes, due to brain maturation (Anderson & Perone, 2018). Results of the linear mixed-effects models indicate that there was no significant effects of linear age or a quadratic age term on children’s neural tracking in any frequency rate or measurement wave. Stressed syllable tracking shows a trend towards a u-shaped pattern, with stronger tracking at 5 months and 2-4 years and a slight decrease of tracking at 10 months, but this did not reach statistical significance in our study (*p* = .06). Therefore, our study does not provide compelling evidence for developmental changes, at least not between infancy and 2-4 years of age when examining SBC in response to sung speech, although we did not sample each age equally thorough. This finding agrees with the results previously reported by Ortiz-Bajaras et al. (2021), who found no major differences in children’s phase tracking of speech between birth and 6 months of age, and their results were numerically similar to those observed in adults (Pefkou et al., 2017). We find no evidence for developmental changes in neural tracking of sung speech from infancy to early childhood.

Apart from more general effects of brain maturation, this also suggests that children’s neural tracking of sung speech, measured with SBC, is not affected by an increase in top-down linguistic knowledge of the native language between 5 months and 2-4 years. Otherwise, we would have found some differences between prelinguistic infants and preschoolers. This suggests that SBC is likely to reflect more exogenous bottom-up processes in response to the acoustic signal. Although previous studies did not report major developmental changes in neural tracking, this does not fully agree with the previous reports by Attaheri, Choisdealbha, et al. (2022) and Attaheri, Panayiotou, et al (2022) who reported small developmental changes. There are at least three possible explanations for the absence of a developmental change in neural tracking. First, a developmental pattern may be obscured by the nature of the stimuli: melodically sung well-known nursery rhymes, for which top-down linguistic processing might be less essential, compared to freely spoken language. A previous study found that adults show increased phase coherence for sung compared to spoken language (Vanden Bosch der Nederlanden et al., 2022). In our stimuli, word onsets were consistently marked by stressed syllables. Therefore, even in older children, word segmentation could still rely strongly on syllabic stress cues for these nursery rhymes. Although children are generally assumed to transition from bottom-up segmentation based on strong syllables to more top-down use of linguistic knowledge and additional cues (e.g., Kidd et al., 2018), this shift may be masked when stressed syllables consistently coincide with word onsets. In addition, increased neural tracking has been observed when songs are perceived as familiar (Vanden Bosch der Nederlanden et al., 2022). Potentially, known nursery rhymes have strong memory representations and highly predictable rhythmic and melodic structure. The increased neural tracking of known songs and more similar top-down predictions across participants can result in reduced individual variability in neural tracking, which can make it harder to detect any small developmental changes. However, the infants in Attaheri, Choisdealbha, et al. (2022) and Attaheri, Panayiotou, et al (2022) also listened to sung or rhythmically chanted nursery rhymes, yet those studies reported small developmental changes using TRFs. This suggests that the discrepancy may not be due to the type of stimulus but rather to differences in the neural tracking measures themselves. Specifically, our specific measure of neural tracking (i.e., SBC) may be less sensitive to developmental changes compared to TRFs. SBC and TRFs may capture slightly different aspects of neural tracking, some of which are more strongly influenced by emerging linguistic knowledge than others.

In line with our main prediction, we found that children’s receptive vocabulary measured at 2-4 years of age was positively associated with the degree of neural tracking, but only in the stress rate regardless of children’s age. This supplements the earlier findings by Menn, Ward, et al. (2022) who found that infants’ neural tracking of nursery rhymes at 10 months – but not 14 months – predicted their vocabulary skills at 24 months. The results of our large-scale replication suggest that neural tracking in the stress rate remains predictive of language development at least until early childhood. Therefore, the hypothesis that the predictive value of neural tracking the acoustics of speech for language development could decrease over time, because children may not need to neurally track the acoustics of speech stream anymore once they have access to more top-down, linguistic knowledge of the language, is not supported by our data. For Dutch, initial word stress is an important cue for speech segmentation (Cutler, 1994). As theorised in the introduction, children attending to word stress in the speech stream might benefit in the word segmentation task, subsequently facilitating language development (Junge et al., 2012; Snijders, 2020). The results suggest that some children show increased sensitivity to stressed syllables or prosodic information in the speech stream, regardless of language experience, which is associated with increases in children’s receptive vocabulary knowledge during preschool years.

While we found the expected positive association between neural tracking in the stress rate and children’s receptive vocabulary outcomes, which can reflect increased attention to stressed syllables and improved word segmentation (Snijders, 2020), there are some alternative explanations. Children with increased SBC in response to sung speech may have been more familiar with the songs compared to other children. Higher familiarity with songs leads to stronger SBC (VandenBosch der Nederlanden et al., 2022). Stronger SBC values could therefore imply that those infant were being sung to more often, which could indirectly have enhanced their vocabulary development (e.g., Torppa et al., 2020). Singing is a highly engaging way to provide language input to infants (Snijders et al., in press). In addition, singing also increases infants’ selective attention to the singers’ mouth which can support language learning (Alviar et al., 2023). Furthermore, the association could also reflect that some infants are more frequently exposed to infant-directed speech than others, resulting in more familiarity and stronger neural responses to this speech register, indirectly supporting their language development (e.g., Weisleder & Fernald, 2013). In addition, more frequent parent-infant interactions supports the development of white matter which is associated with language outcomes (Huber et al., 2023), potentially increasing how efficiently neural responses lock onto external stimuli. In future studies, it can therefore be insightful to examine how the amount of infant-directed speech and song in the infant’s environment is related to their neural responses to this speech register; and subsequently, how these measures uniquely predict variation in children’s language development.

In conclusion, the present study examined SBC in a uniquely large longitudinal sample including over 2000 children. The children listened to Dutch sung nursery rhymes while EEG was being recorded at 5 months, 10 months, and 2-4 years. First, we found that infants from 5 months onwards neurally track the frequencies of stressed syllables, syllables, and phonemes in Dutch nursery rhymes. Unexpectedly, there were no developmental changes in neural tracking across different frequency bands from prelinguistic infants into early childhood. SBC in the studied frequency bands likely reflects exogenous bottom-up processes in response to the acoustic signal. Second, and in line with our expectations, we found that children with larger receptive vocabularies at 2-4 years showed increased neural tracking in the stress rate. The predictive value of stressed syllable tracking did not change across development. The findings that stressed syllable tracking does not change between 5 months and 2-4 years, and that the degree of stressed syllable tracking is positively related to children’s receptive vocabulary outcomes at 2-4 years, suggests that some children show more attention to stressed syllables, which is related to their language skills from infancy to preschool years.

## Data availability

YOUth is a longitudinal study that aims to produce and safely store FAIR and high-quality data. The data can be accessed for both use and verification purposes upon request (see https://www.uu.nl/en/research/youth-cohort-study/data-access). The preregistration, *R* script, and other materials can be found online on OSF: https://osf.io/pq6e5/

## Acknowledgements

ChatGPT was used for minor language editing and figure formatting. All content, analysis, and conclusions are the work of the authors.

We would warmly like to thank Ádám Takács, Adhyayan Chaudhry, and Marlene van Lierop for their help in designing and programming the automated EEG preprocessing pipeline (see Chaudhry et al., 2025).

